# Sea Surface Temperatures Drive Historical Demography of Deep-Sea Fishes

**DOI:** 10.1101/2021.07.07.451542

**Authors:** Max D. Weber, Joshua E. Carter, Ron I. Eytan

## Abstract

Demographic histories are largely understood to be a product of their environment, as populations expand or contract in response to major environmental changes. Deep-pelagic fishes inhabit one of the most temporally and spatially stable habitats on the planet, so they may be resistant to the demographic instability commonly reported in other marine habitats, but their demographic histories are poorly understood. We reconstructed the demographic histories of thirteen species of deep-pelagic fishes using mitochondrial and nuclear DNA sequence data. We uncovered widespread evidence of demographic expansion in our study species, a counterintuitive result bases on the nature of the deep-pelagic. The frequency-based methods detected potential demographic changes in eleven species, while the Extended Bayesian Skyline Plots were more conservative and identified population expansion in five species. The dates of expansion largely coincide with periods of warm sea-surface temperature at the northern and southern boundaries for the ranges these species inhabit. We suggest that this is the result of the pelagic larval phase shared by most deep-pelagic fishes, where the larvae inhabit the upper 200 meters. Changes in sea surface conditions likely alter the suitability of the habitat in a given region for the larval phase, affecting the species range and in turn population size. These results are critical to our understanding of how the deep-pelagic fish community will respond to future climatic changes.

## INTRODUCTION

The demographic history of a species is strongly influenced by the environment it inhabits (Alheit & Hagen 1997; Avise et al. 2000; Grant 2015). Major changes in the environment can alter the distribution and size of suitable habitat for a species, reducing or increasing the species’ range. Population sizes expand or contract in response to these fluctuations in range size (Avise et al. 2000; Nye et al. 2009). Evidence for the environment’s control over population dynamics can be seen across taxonomic groups in terrestrial and marine habitats around the world (Almada at al. 2012; Eytan & Hellberg 2010; Grant 2015, Robalo et al. 2012).

Given that changes in environmental conditions strongly influence population size, species inhabiting unstable environments should be characterized by unstable population sizes. On the other hand, species inhabiting temporally stable environments should be less susceptible to frequent population expansions or contractions due to global climatic events. Studies have supported this notion, finding genetic diversity to be greater in species inhabiting more stable environments than closely related species in environments more subject to change (Carnaval et al. 2009; Gugger et al. 2013).

The deep-pelagic, the marine habitat found between 200 meters in depth to approximately 100 meters above the sea floor, is considered one of the most temporally and spatially stable environments on the planet. In terms of physical characteristics like temperature, there is a strong latitudinal homogeneity in the environment that increases with depth (Robison 2009). In comparison to shallower habitats, temperature change occurs slowly over time and the magnitude of change is less (Abraham et al. 2013; Clark et al. 2006; Clark et al. 2009; Levitus 2000; Levitus 2012; Mora et al. 2013; Robison 2009). Based on the stability of the environment and our current understanding of the manner in which habitat influences demography, the population sizes of the fishes inhabiting the deep-pelagic would be expected to be stable over time.

If the demographic histories of deep-pelagic fishes are not stable, it is difficult to predict which physical factors could drive this instability. Insights into the historic demography of deep-sea organisms are few, and have focused on deep-benthic species (Etter et al. 2005; Sakuma et al. 2014; Varela et al. 2012). The deep-benthic environment, benthic habitat found below 200 meters, is much more heterogeneous than the deep-pelagic and likely under differing environmental pressures (Thurber et al. 2014).

The physical factors influencing the historical demography of marine fishes inhabiting the less stable upper 200 meters are better understood. Studies have consistently shown population size changes that correspond with major changes in the environment that would have far less dramatic effects on the deep-pelagic. A plurality of these studies report widespread population expansion in shallow-dwelling fishes following the last glacial maximum. Two factors are frequently cited to explain increases in geographic range and a corresponding increase in population size: an increase in global sea-surface temperatures and sea-level rise that dramatically increased in shelf habitats (Avise et al. 2000; Eytan & Hellberg 2010; Grant 2015). During this same period there would have been a significantly less increase in temperatures in the deep-pelagic and the amount of deep-pelagic habitat increased negligibly (Abraham et al. 2013; Clark et al. 2006; Clark et al. 2009; Levitus 2000; Levitus 2012; Mora et al. 2013; Robison 2009).

If the demographic histories of deep-pelagic fishes are unstable it is more likely that physical conditions outside of the deep-pelagic drive these dynamics. Two features of deep-pelagic fishes could be responsible: diel vertical migration and a pelagic larval phase. Many deep-pelagic fish species perform diel vertical migrations, a daily migration to the shallower and more variable epipelagic waters (Barham 1966; Sutton 2013). An analysis of the distribution of deep-pelagic fishes found that the ranges of vertically migrating species were more likely to change in response to large scale changes in climate than the ranges of species that do not vertically migrate (Hsieh et al. 2009). This could be a result of significantly greater heating in the upper ocean than deep waters, which vertical migrators visit on a daily basis. If the changes in surface waters are no longer tolerable to the vertically migrating fishes these species can no longer persist in their former range. If vertical migratory behavior drives demography in deep-pelagic fishes, vertical migrators will be characterized by population expansions and/or contractions, while the population size of non-vertical migrators will be relatively stable over time.

Yet another explanation for unstable deep-pelagic fish populations could be the existence of a pelagic larval phase in nearly all deep-pelagic fishes (Bowlin 2016; Johnson et al. 2009; Moser 1996). These larvae reside in the upper 200 m and are subjected to conditions that are more variable than those experienced by the adult forms. Changing conditions in surface waters could alter larval distribution and in turn affect the species range and population sizes.

Knowledge of the demographic history of deep-pelagic fishes serves two key purposes. First, it will provide insight into the ecological processes driving population dynamics in one of the world’s most stable environments. This could help to bolster current theory that suggests that such a stable environment should produce stable populations. Secondly, understanding how deep-pelagic fishes responded to past climatic events will allow us to make predictions about how they will respond to future changes in climate. This is critical given the ecological importance of midwater fishes. The deep-pelagic comprises approximately 95% of the ocean by volume, and the biomass of deep-pelagic fishes is at minimum ~1,000 million tons, several orders of magnitude larger than the total global commercial fisheries landings (Gjosaeter & Kawaguchi 1980; Irigoien et al. 2014). Furthermore, deep-pelagic fishes are important prey items for numerous commercially targeted species (Battaglia 2013; Varghese et al. 2013).

To better understand the demography of deep-pelagic fishes, we answered the following questions: (1) Are deep-pelagic fish populations stable over time? (2) If population expansions or bottlenecks are detected, can the results be explained by adult migratory habits or (3) the pelagic larval phase in deep-pelagic fishes?

## METHODS

### Sampling and Sequence Generation

We selected thirteen deep-pelagic species that span taxonomic groups, life histories, and vertical migration behavior (See Supplementary Table 2). Samples were obtained by trawling with a MOCNESS net system in depth zones up to 1,500 meters in the northern Gulf of Mexico (Cook et al. 2020) (Figure 1). Upon collection and identification of vouchers at sea, a ~1cm piece of lateral muscle tissue was preserved in ethanol. Voucher specimens are housed in the Ocean Ecology Lab at Nova Southeastern University.

**Figure 1.**
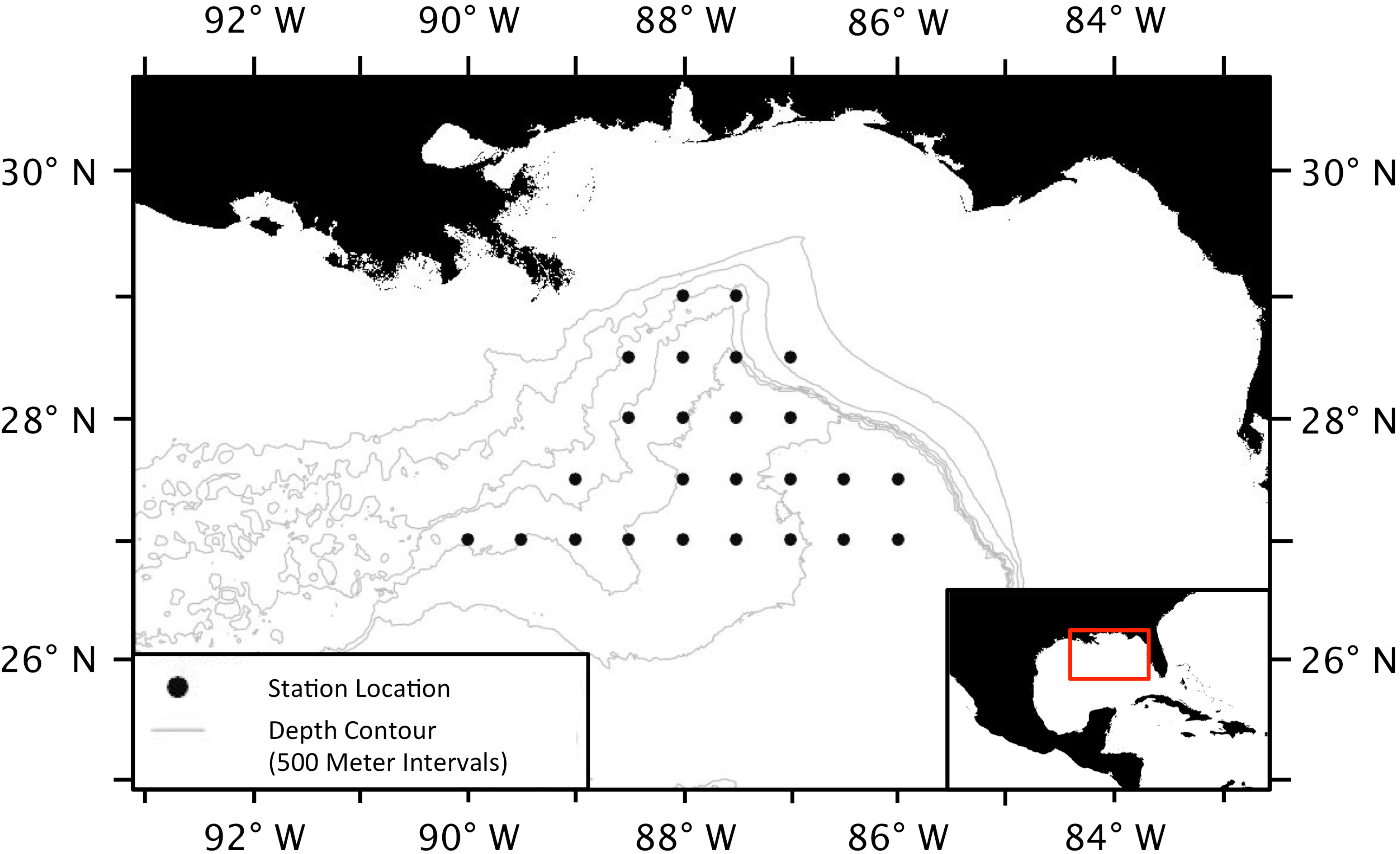
Map of the study area. Samples were collected in the Northern Gulf of Mexico. Black dots indicate stations where sampling took place.

We generated DNA sequence data from the mitochondrial gene *cytochrome oxidase* I (*COI*) as well as three nuclear DNA exons (PLAG, ENC, and MYH). PCR was performed using Promega GoTAQ (Madison, WI) (see Supplementary table for primers used). Following amplification, all PCR products were cleaned using a standard PEG protocol (Glenn 2019). Amplicons were Sanger-sequenced on an ABI 3730 capillary sequencer at Yale Keck Biotechnology Resource Laboratory. Sequences were cleaned and edited in Sequencher v5.1. Nuclear markers were phased using Phase v2.1 to resolve heterozygous sites. The sequences were then aligned using MAFFT in Generiousv9.1.8 (Kearse et al. 2012).

### Frequency-Based Analyses

We calculated Tajima’s D, Fu’s Fs, and R2 for each marker (Fu 1997; Ramos-Onsins & Rozas 2002; Tajima 1989). Comparisons of the statistical power of frequency-based tests have shown that Fsand R2are the most capable of detecting population growth (Ramos-Onsins & Rozas 2002). They complement one another as well, with Fss excelling at population growth detection in large sample sizes, while R2 2 performs better with small sample sizes. A significant and large negative Fs value suggests population growth, while a significant and small positive R2 value indicates population growth. Tajimas’s *D* points to population growth and/or a selective sweep when significant and negative.

All of the frequency-based tests were performed in DNAsp v6 (Rozas et al. 2017). Ambiguity codes were replaced with Ns to allow for calculation in DNAsp. Tajima’s *D* is a two-tailed test, so significance was initially determined by the test itself. The significance of all three tests was also determined using coalescent simulations with 1000 replicates implemented in DNAsp.

### Gene Tree Based Analysis

The second set of tests makes use of the topologies and branch lengths of gene trees to infer changes in population size over time using the coalescent. We performed these analyses in BEAST v2.4.7 (Bouckhaert et al. 2014) to generate Extended Bayesian skyline plots (EBSPs). EBSPs utilize coalescent theory and a Markov Chain Monte Carlo Algorithm to infer and visualize demographic changes in a dataset. The Bayesian skyline plot is preferable to earlier skyline plot methods as it models both genealogy and demographic history simultaneously, which reduces error rates from uncertainty in estimates of node time (Heled & Drummond 2008; Ho & Shapiro 2011).

Nuclear and mitochondrial genes were included in the analysis for each species. The chain length was set to 50,000,000 sampling every 1,000. Initial clock rates were derived from secondarily calibrated ultrametric trees based on the results of Near et al. 2013. A strict clock was set and the clock rate of COI was fixed. The nuclear rates were given a normal distribution with a median set to the initial clock rate. The partitioning scheme and substitution models were set based on Partition Finder v2.0 results (Lanfear et al. 2012).

A second set of trees was created using the same methodology, with the exception of the selection of substitution models. All partitions were set to the RBS substitution mode (Bouckaert et al. 2014). RBS is a reversible-jump based substitution model for nucleotide data. This substitution model does not require a fixed substitution model to be assigned to each partition at the beginning of the analysis. Instead, it allows five different substitution models to be explored through the run, in order to find the substitution model with the best fit to the dataset.

After running in BEAST, log files for both sets of trees were inspected using Tracer v 1.7.1 (Rambaut et al. 2018). The most strongly supported EBSP analysis, based on ESS values, was selected and used for the inference of each species’ demographic history. The posterior estimate of the number of population size changes provided a test for a rejection of constant population size. Finally, the trees files were uploaded to Rstudio v 0.99.484 (Studio 2012). The Rscript “plotEBSP”, provided with the EBSP tutorial (http://www.beast2.org/files/2016/01/ebsp2-tut.zip), was used to generate and visualize the extended Bayesian skyline for the timing of inferred population size change events (Heled 2010). Vertical lines were added to the plots to indicate the approximate timing of the onset of population expansion.

### Population Dynamics and Vertical Migration

We placed species into two categories; those that had undergone an inferred population size change and those that had not. Species placed into the “inferred population size change” group were categorized as such if the inference was uncovered in both the frequency-based and gene tree based analyses. We further divided species into vertical migrators and non-vertical migrators. A chi-squared test was used to test for a correlation between inferred population size changes and vertical migration. Information on the migration patterns of two fishes (*Polymixia lowei*, and *Synagrops spinosu*s) was unavailable, so they were left out of the analysis.

### Sea Surface Temperature and Population Size Changes

We plotted sea surface temperatures (SST) for the past 300,000 years from the North and South Atlantic based on previously published climate data (Becquey & Gersonde 2003; Ruddiman et al. 1989). We selected these datasets as they provided sufficient resolution/coverage for the period of time when inferred demographic changes occurred. Furthermore, the two datasets were obtained from sites located approximately 41**°**N and 42**°**S, corresponding to the northern and southern extent of the range inhabited by many tropical/low-latitude deep-pelagic species. Accordingly, changes in SST at these latitudes could directly impact the larval distribution for low-latitude deep-pelagic species. We indicated the mean temperature, as well as the mean temperature plus one standard deviation for this period, to identify periods that were warmer than average as well as period of exceptional warmth. The dates of the onset of population size change inferred from our EBSPs were mapped on the plot to allow for the comparison of fluctuations in SST and population dynamics in our study species.

## RESULTS

### Summary

The number of sequences generated for each species and marker varied according to sample availability and our ability to achieve amplification. The number of unique sequences obtained for each gene ranged from a low of ten to a high of 97. We used two genes for analysis in ten species, three genes in one species, and four genes in two species (Table 1). Sequences have been deposited to GenBank with #s xxxxx-xxxxx.

**Table 1.**
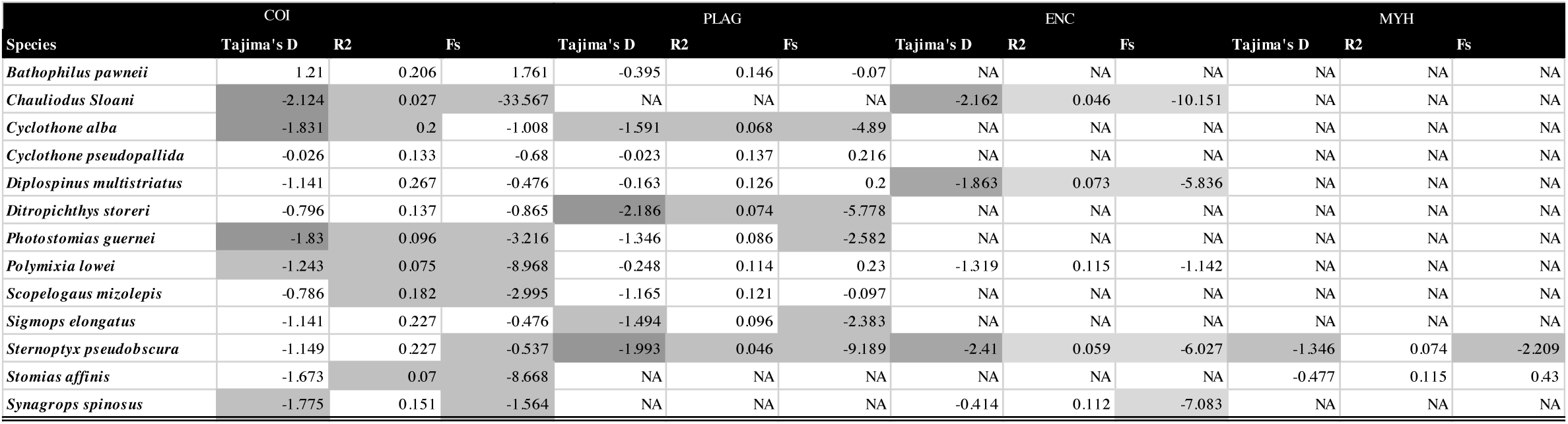
Results of frequency-based statistics analysis. Tajima’s D values that were significant based on the two-tailed test are dark grey. Significant values determined through coalescent simulations are highlighted in light grey.

### Frequency-based analyses

Frequency-based analyses suggested population expansions in 11 of our 13 sampled species (Table 1). In four species (*Chauliodus sloani*, *Sternoptyx pseudobscura*, *Cyclothone alba*, and *Photostomias guernei*) more than half of the frequency-based test for all markers suggested population size changes. Weaker support was present in another seven species (*Diplospinus multistriatus, Ditropichthys storeri, Polymixia lowei, Scopelogaudus mizolepis, Sigmops elongatus, Stomias affinis,* and *Synagrops spinosus),* where less than half of the markers tested produced significant results. No evidence for demographic change was present in *Bathophilus pawneii* or *Cyclothone pseudopallida*.

### Gene Tree Based Analysis

We were able to reject a constant population size in five of the thirteen species (Figure 2). We inferred increases in population sizes for all of the species where a constant population size was rejected. The timing of these population expansions was highly variable. *Cyclothone alba* experienced the most recent increases in population size, which appears to have begun about 85 Kya. Three other species experienced population size increases 125-200 Kya (*Chauliodus sloani*, *Photostomias guernei, and Polymixia lowei*). The oldest population increase that we inferred occurred approximately 270 Kya in *Sternoptyx pseudobscura*.

**Figure 2.**
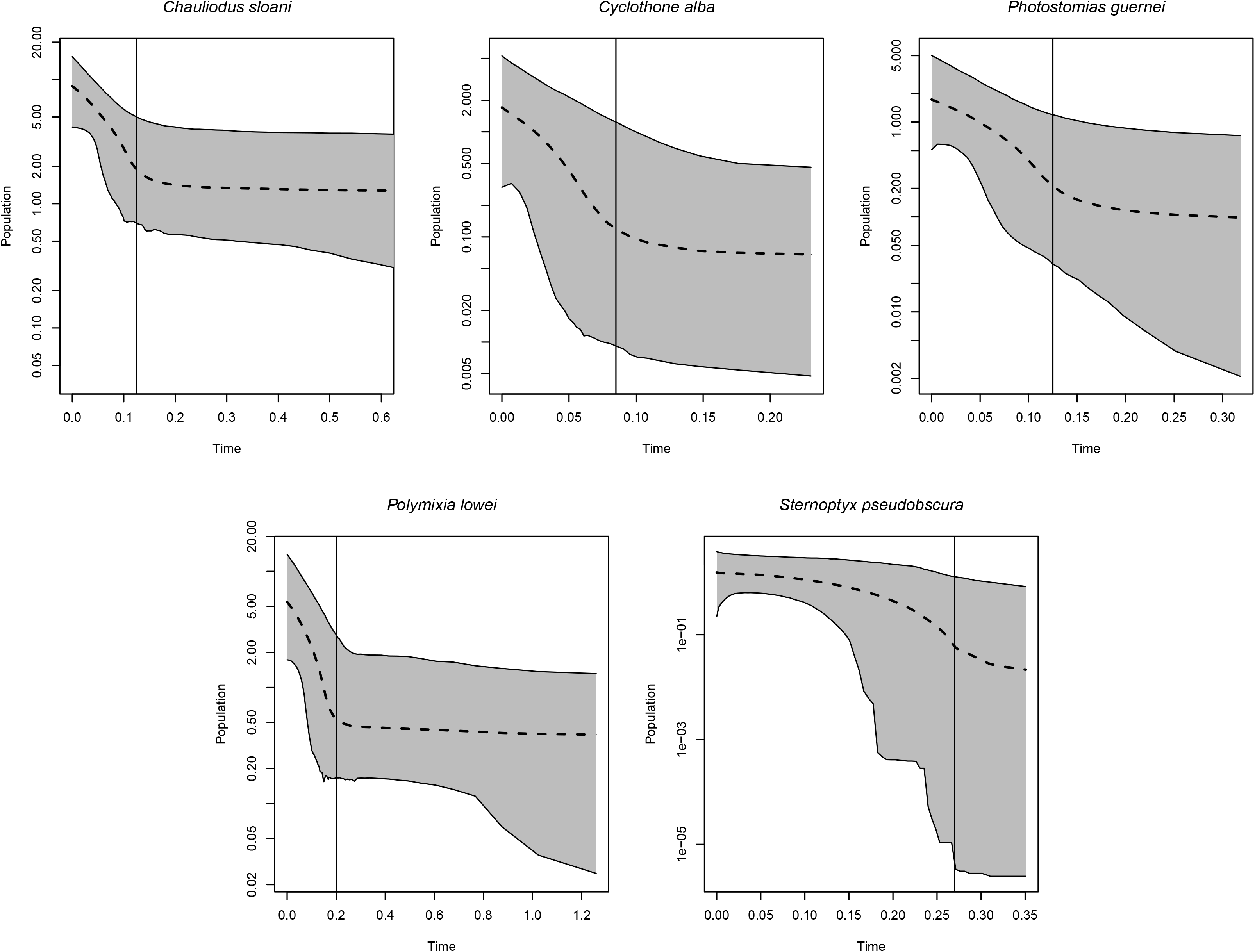
Extended Bayesian Skyline Plots. The x-axis represents time scaled in terms of millions of years, while the y-axis represents relative population size on a log scale. The grey area displays the 95% central posterior density. The vertical black line indicates the approximate onset of population expansion.

### Frequency-based Analyses vs EBSPs

Frequency-based tests suggested population expansions in eleven species. The four species with the strongest support in the frequency-based tests were all found to have undergone expansions using the EBSP method. Our EBSP analyses also recovered a population expansion in *Polymixia lowei*, which had moderate support (fewer than half of the tests were significant) in the frequency-based analyses.

### Vertical Migration and Population Dynamics

The chi-squared test did not provide support for a relationship between vertical migration and population size changes (p-value 0.8190, Table 2).

**Table 2.**
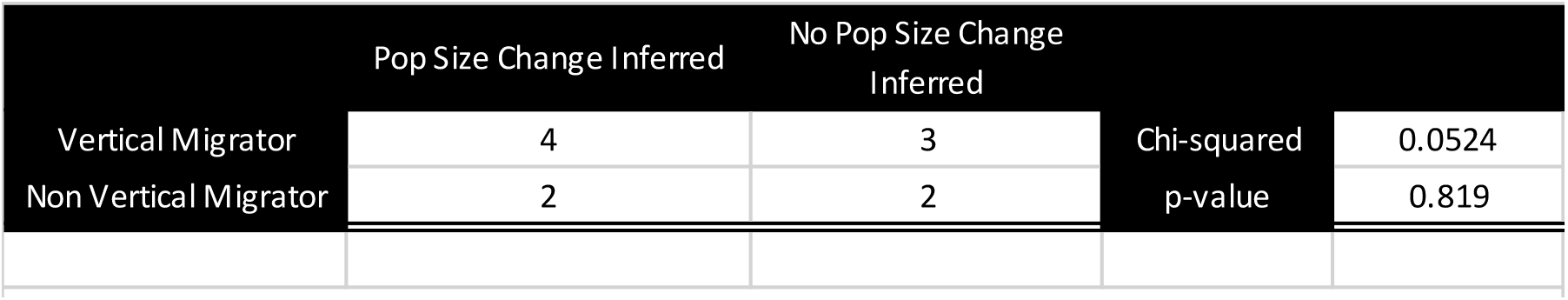
Chi-squared test for significance of vertical migration on the inference of recent population size changes.

### Sea Surface Temperature and Population Size Changes

In our analysis, there were thirteen instances when the reconstructed SST in the North Atlantic reached the mean temperature plus one standard deviation (Figure 3). The reconstructed SST in the South Atlantic reached the mean temperature plus one standard deviation nine times. The general warming and cooling trends were largely in agreement between the two sites.

**Figure 3.**
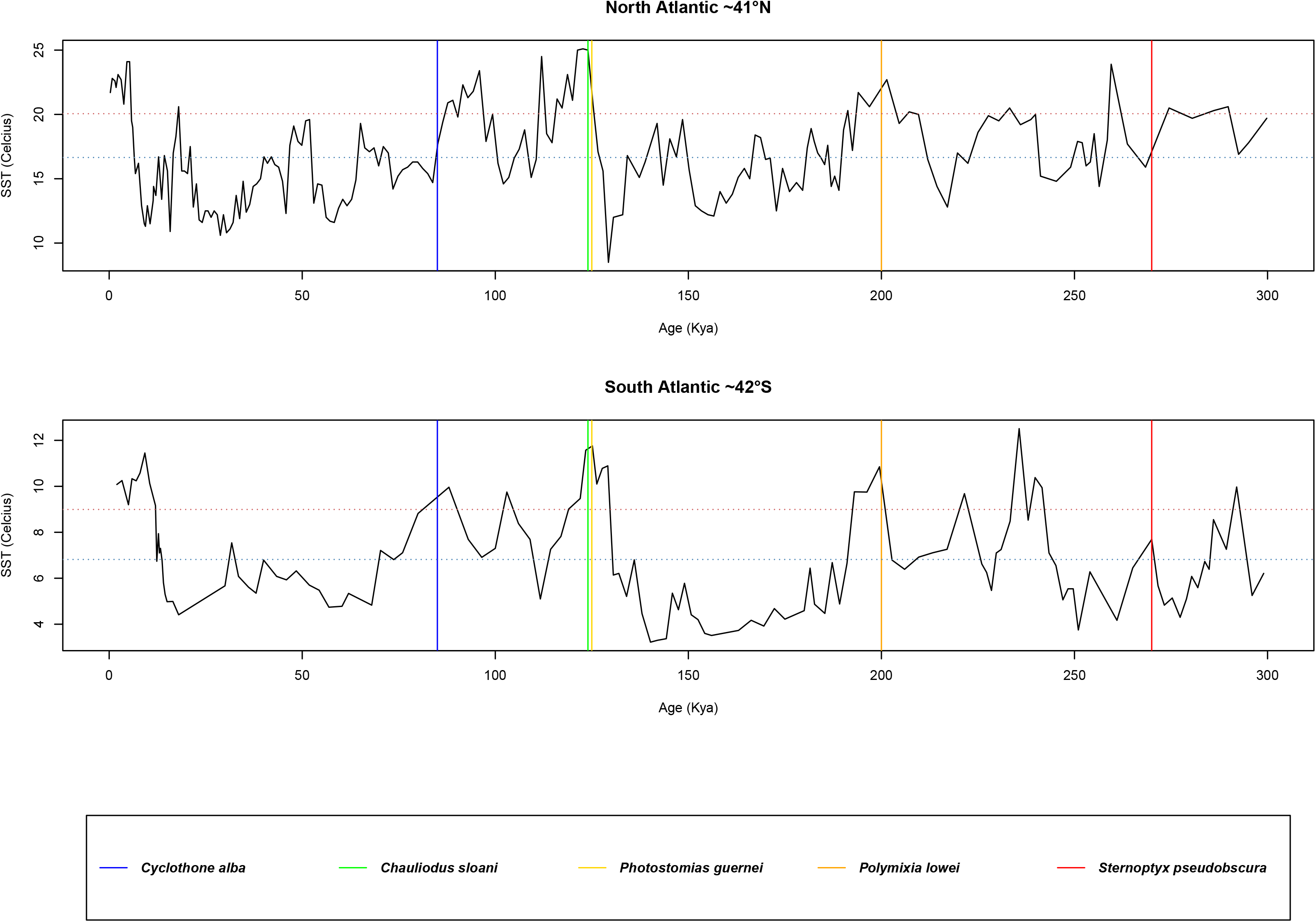
SST in the North and South Atlantic in relation to the onset of population expansion. The x-axis represents time scaled in terms of millions of years, while the y-axis represents temperature in degrees Celsius. The horizontal blue dotted line indicates the mean temperature during this period of time, and the horizontal red dotted line indicates the mean temperature plus one standard deviation. Vertical lines show the onset of population expansion for each species.

We plotted the approximate onset of expansion for all five species that underwent demographic expansion according to our Extended Bayesian Skyline Plot analyses. The population expansions for *Chauliodus sloani*, *Photostomias guernei*, and *Polymixia lowei* all begin during periods of exceptionally warm SST at both sites, where temperatures greatly exceeded the mean plus one standard deviation. *Cyclothone alba* commences expansion during a warm period where the South Atlantic temperature exceeded the mean plus one standard deviation, however it falls just outside of a similar warm period in the North Atlantic. *Sternoptyx pseudboscura* is an outlier in this regard, and appears to take place under conditions where the SST is close to the mean for this three-hundred-thousand-year period.

## DISCUSSION

Historic changes in effective population size are frequently inferred for marine fishes and attributed to major ecological events and past climatic change (Avise et al. 2000; Grant 2015). Previous studies however, have largely focused on species inhabiting shallower environments that are more variable over time in terms of physical conditions such as temperature in comparison to the deep-pelagic. Given the stability of the deep-pelagic, populations of fishes inhabiting this environment might also be stable, so that no effective population size changes would be inferred from genetic examination. Nonetheless, we uncovered multiple lines of evidence suggesting population expansions in five of our thirteen deep-pelagic study species, the most ancient event occurring about 270 Kya. These expansions and their timings were based on our EBSPs, which utilized both mitochondrial and nuclear data. Four of the five dates of population expansion coincide with periods of warm sea surface temperatures (SST) at 41° North and 42° South in the Atlantic, a transition zone for many tropical deep-pelagic species. Future changes to SST in this region due to climate change would impact the distribution of deep-pelagic larvae residing in surface waters, impacting deep-pelagic fish population sizes.

### Both Frequency-Based Statistics and EBSPs Recover Evidence for Population Expansions

Population expansions were recovered using two different types of analyses that were largely in agreement, however the EBSPs suggested fewer instances of demographic change. It is not surprising that these tests would not agree in every case. Simulated datasets demonstrate that the EBSP analyses are prone to false negatives when using fewer than eight loci (Heled & Drummond 2008). Given the number of loci sequenced it seems likely that our EBSPs were more conservative in their inference of population expansions than the frequency-based tests.

### Deep-Pelagic Fish Population Dynamics Likely Shaped by Pelagic Larval Phase

Our widespread inference of population expansions in deep-pelagic fishes was unexpected. Demographic events are typically attributed to major environmental changes that alter the environment, in turn increasing or decreasing the amount of suitable habitat for a given species (Avise et al. 2000; Grant 2015). Populations expand or contract in response to these alterations in available habitat. Because the deep-pelagic has been a relatively stable habitat in terms of its size and temperature for millions of years, it would seem likely that the demographic histories of deep-pelagic fishes would be characterized by a lack of expansions/contractions (Clark et al. 2009; Levitus et al. 2012; Robison 2009). Instead, we uncovered a minimum of five cases of population expansion (identified by both frequency-based statistics and gene tree based analysis) within the past 300,000 years.

We proposed two potential drivers of population size change in deep-pelagic fishes: vertical migratory habits and the pelagic larval phase. Both hypotheses were based on obligate use of more physically variable surface waters by the study species. Unlike the deep-pelagic, surface waters are characterized by both frequent and dramatic changes in conditions such as sea surface temperature (SST). While the conditions at depth may remain favorable to a species, surface waters conditions could impact species distribution.

Our first proposed driver of demographic change focused on diel vertical migration (DVM), the daily migration to surface water by deep-pelagic fishes. Hsieh et al. (2009) provides support for this hypothesis as they reported the larval distribution of fish species with vertically migrating adults changed more rapidly/frequently than non-vertically migrating species. This could be attributed to short-term changes in surface water conditions that impact vertically migrating species but are unfelt by those adults that remain at depth. If vertical migratory habits were the primary driver of population dynamics in deep-pelagic fishes, the demographic histories of vertically migrating species would be characterized by population expansions/contractions, while non-vertically migrating species should be stable over time. Based on our chi-squared test we were unable to detect any difference in population dynamics between these two groups, and reject this hypothesis.

The second mechanism we investigated was larval control over the demographic histories of deep-pelagic fishes, as the larvae of most deep-pelagic fishes reside in the upper 200 meters (Bowlin 2016; Johnson et al. 2009; Moser 1996). Two lines of evidence support this hypothesis: long-term monitoring of larval distribution and deep-pelagic patterns of distribution.

Long-term monitoring efforts in transition zones between tropical and subpolar regions have shown that physical conditions, such as sea surface temperature, are key predictors of larval community composition. Furthermore, physical changes in these environments alter the larval composition of the community (Ahlstrom 1969; Netburn & Koslow 2018; Sassa et al. 2004; Urias-Levya et al. 2018). Aceves-Medina et al. (2004) found that the distribution of larvae was congruent with that of the adults. This suggests that as sea surface conditions alter larval distributions, the ranges of adults would change accordingly.

The second piece of evidence for larval control on demography comes from the distribution patterns of deep-pelagic fishes. Deep-pelagic fishes can broadly be classified as tropical or polar species, and tend to have clear latitudinal geographic boundaries to their ranges (See Supplementary Table 3 for range description of study species) (Olson 2001; Pearcy 1991; Randall 1981). Within oceanic basins, latitudinal differences in temperature decrease by depth. By 1000 meters the temperature is a near uniform 5 degrees Celsius throughout most of the world’s oceans (Helfman et al. 2009; Tyus 2011). It is therefore noteworthy that even the range of non-vertically migrating bathypelagic groups such as the whale fishes fit this pattern (Paxton 1989). It seems unlikely that this distribution can be explained by physiological constraints on adults of these species given the relative homogeneity of the environment. Rather, a given species range is constrained to regions with surface waters tolerable to their larvae. If correct, we hypothesized that periods of warm SST in high latitudes would increase the range of tropical deep-pelagic fishes, and lead to population expansions.

To explore this hypothesis we compared our dates of inferred expansions to reconstructed SSTs for the Atlantic at 41° North and 42° South. We inferred five cases of population expansion with our EBSPs, and all but one appear to coincide with warm periods of SST at this site, the northern and southernmost boundary for many tropical deep-pelagic fish ranges (Olson 2001). This supports the notion that sea surface conditions constrain the ranges of species living far below them through their larval phase, and that they have profound impacts on their population sizes. If the tolerances of pelagic larvae dictate species distribution, it would explain why we were unable to detect any difference between vertically migrating and non-vertically migrating adults. The SST conditions present in a region will lead to a pelagic larval community that will mature into adults tolerant to those conditions. In this manner, the physiological tolerances of pelagic larvae seem to drive the demographic histories of deep-pelagic fishes.

Another interesting result was the frequency-based performance of *Cyclothone pseudopallida* and *Sigmops elongatus*. These species had very little to no support for population expansions across all genes. While our other study species have clearly defined low latitude distributions, these two species exist well into polar waters in the northern hemisphere. It would seem that their larvae have much greater tolerances for a wide range of conditions. Our hypothesis predicts that these species would be less susceptible to expansions/contractions in the face of climate change and our analysis found this to be the case.

### Future Directions

Exploration of the demographic histories of polar mesopelagic and bathypelagic fishes could provide further support for our findings. If periods of warm sea surface temperatures benefit low latitude species, polar species would be expected to experience population bottlenecks during these times. Instead, high-latitude species would only undergo range expansion and a resultant population expansion when sea surface temperatures were low. Repeating this methodology on a set of polar deep-pelagic fish species could demonstrate whether this is the case.

## Conclusions

Insights into the mechanisms that control deep pelagic-fish population dynamics are lacking. Our results demonstrate that despite the long-term stability of the deep-pelagic, the population sizes of the fishes that reside in this habitat are not static in nature. The dates of expansion we inferred suggest that low latitude deep-pelagic fish species respond positively to an increase in sea surface temperature at high latitudes. As we come to understand the environmental factors that influence demographic changes in these fishes we will better be able to predict how populations of these fishes will behave in the face of future climate change.

